# N-terminal acetylation of superoxide dismutase 1 accelerates amyloid formation without general destabilization of the apo-state

**DOI:** 10.1101/2025.03.28.645989

**Authors:** Kristine Steen Jensen

## Abstract

Co- and posttranslational modifications can significantly impact the structure, dynamics and function of proteins. In this study we investigate how N-terminal acetylation affects misfolding and self-assembly of the enzyme superoxide dismutase 1 (SOD1), implicated in amyotrophic lateral sclerosis (ALS). Studies of protein inclusions in patient samples and animal models have shown that wild-type SOD1 can form amyloid fibrils even when no mutations are found in the *sod1*-gene. This has identified SOD1 amyloid formation as a possible common denominator of ALS and may suggest that co- and posttranslational modifications, like N-terminal acetylation found in human SOD1, can be a factor in disease development. In this work the impact of N-terminal acetylation of SOD1 on stability and aggregation is characterized. Results show that the structure and thermal stability of the apo-state are unaffected by the modification while the amyloid formation rate is significantly enhanced. This is caused by a shortening of the nucleation phase together with an increase of fibril elongation by more than 10-fold upon N-terminal acetylation of SOD1. Collectively the findings demonstrate how regulation by co- and posttranslational modifications can influence protein misfolding and self-assembly.

## Introduction

Protein misfolding and amyloid formation is a hallmark of many human neurodegenerative diseases including Alzheimer’s- and Parkinson’s disease, diabetes and amyotrophic lateral sclerosis (ALS). Most of these diseases have middle to late-life onset although in multiple cases the disease is coupled to specific inhered gene mutations present from the onset of life in all cells of the body. The molecular triggering points and disease mechanisms are in many cases not well understood and to date only very limited treatments for proteinopathies are available (1).

ALS is a fatal neurodegenerative disease leading to motor neuron death and central nervous system dysfunction. ALS has a broad spectrum of phenotypes, and variations in many genes are coupled to an increased risk of ALS (2, 3). More than 200 different single-point mutations in the gene encoding the cytosolic enzyme superoxide dismutase 1 (SOD1) are associated with familial forms of ALS and lead to a toxic gain of function. Investigations of protein inclusions in nerve and brain tissues from individuals who suffered from sporadic ALS or familial ALS caused by mutations in other genes than *sod1* have shown wild-type SOD1 to form inclusions in spinal and cortical motor neurons (4, 5). This implicates SOD1 misfolding and amyloid formation as a possible common denominator of ALS. Most cases of ALS are sporadic (estimated to correspond to 90-95% of all reported cases), meaning that there is no known family history of ALS.

SOD1 is a cytosolic homo-dimeric enzyme expressed in most human cells. The active enzyme has an intermolecular disulfide bond between C57 and C146 in each subunit and binds one Zn^2+^ and one catalytic Cu^+^/Cu^2+^ ion per subunit. SOD1 acts as a radical scavenger catalyzing the dismutation of two superoxide radicals (O_2_^·-^) to molecular oxygen (O_2_) and hydrogen peroxide (H_2_O_2_) per catalytic cycle (6, 7).

Posttranslational modifications (PTMs) of proteins are important in the regulation of cellular function. A vast number of different covalent modifications take place in eukaryotic cells enabling reversible regulation of function or targeting of proteins for specific localization or degradation (8). A subset of common PTMs involves small covalent modifications such as phosphorylation, nitrosylation, methylation and acetylation. These small modifications are in some regards comparable to single point mutations; they create new functionalities of the modified sidechains and may exhibit large effects on the structure and dynamics of the modified protein. In addition to sidechains, the N-terminus of proteins are often processed and modified (9). N-terminal acetylation is a common co-translational modification of proteins occurring while the nascent chain is still bound to the ribosome (figure 1) (8). Mass-spectrometry-based proteomics has demonstrated that a large percentage of human proteins are N-terminally acetylated (10). N-terminal acetylation of SOD1 from human erythrocytes has been shown (11), and similar results were found in a large proteomic analysis (10). However, the biological function of N-terminal acetylation of SOD1 is largely unknown and little is understood about how the physico-chemical properties of SOD1 changes upon N-terminal acetylation.

**Figure 1.**
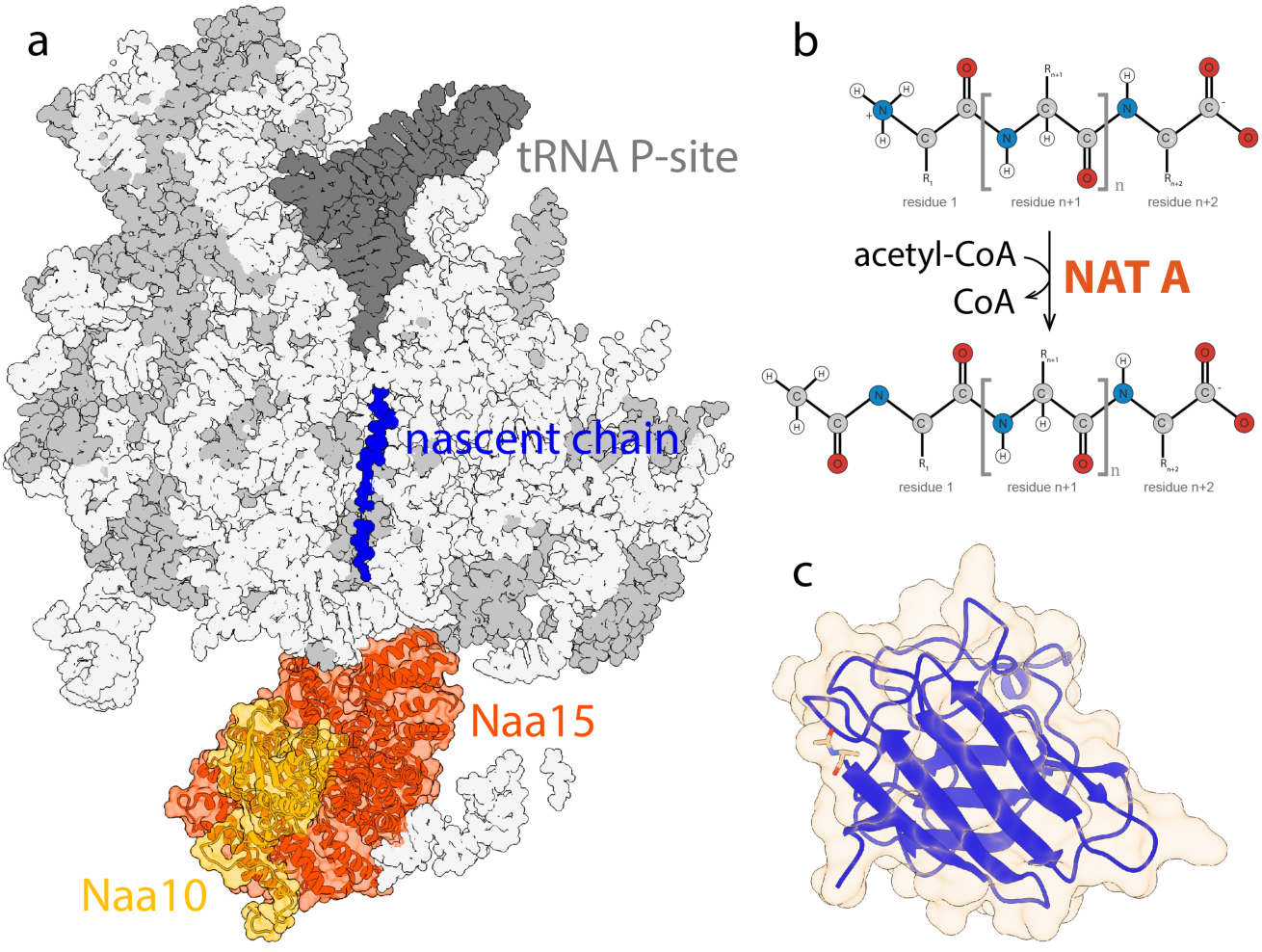
N-terminal acetylation. a) Slice through translating ribosome based on the structure of human ribosome-NAT A complex (PDB entry: 6hd7). N-terminal acetylation is catalyzed by N-terminal acetyltransferase complex on the growing nascent chain after the co-translational action of aminopeptidase to remove the N-terminal initiator methionine residue. We co-expressed the Naa10 (yellow) and Naa15 (orange) subunits of the NAT A complex together with human pwtSOD1^Δcys^ (blue) in E. coli. Naa10 and Naa15 acted together with endogenous ribosomal proteins (gray) and ribosomal RNA (light gray) to produce N-terminal acetylated pwtSOD1^Δcys^. tRNA in the P-site are illustrated in dark gray. b) Upon acetylation of the N-terminal amino group by NAT A the charge of the N-terminal at physiological pH changes from +1 to neutral. C) SOD1 monomer (PDB entry: 2xjl) with modified N-terminal showing the position of the acetyl-group added upon N-terminal acetylation of SOD1.

In this study, we recombinantly express N-terminal acetylated SOD1. N-terminal acetylation is catalyzed by a group of enzyme complexes named N-terminal acetyltransferases (9, 12). This group contains six different complexes named NAT A to NAT F. The different NAT complexes have different substrate specificity, which is determined by the first two amino acids in the polypeptide chain of the substrate protein (12, 13). NAT A is the N-terminal acetyltransferase complex with the broadest substrate specificity acting on polypeptides, starting with Serine, Alanine, Threonine, Glycine, and Valine at the first position adjacent to the translational-initiator methionine. N-terminal acetylation takes place after cleavage by methionine aminopeptidase of the N-terminal translational-initiator methionine residue. The NAT A complex consist of a catalytic subunit NAA10, a ribosome anchoring subunit NAA15, as well as the huntingtin-interacting protein K (HYPK) and NAA50 subunit (9). Only the NAA10 and NAA15 subunits are needed for the enzyme complex to form and be active inside the cytosol of *E. coli* during heterologous expression (14).

Acetylation of the free amino group leads to a change in size and charge of the N-terminus. At physiological conditions with a pH near 7, proteins typically have a positively charged N-terminus. N-terminal acetylation leads to a change of charge from +1 to neutral at physiological pH (figure 1b). The consequence is a reduction of the local charge and a more negative net charge of the protein. It also introduces an additional methyl group with an increase in hydrophobicity. Taken together, these changes may appear minor. However, ongoing work points to the fact that changes in cellular regulation of N-terminal acetylation are linked to multiple disease-causing events (12). However, the understanding of *how* and *why* specific changes in N-terminal acetylation status affect protein function is still limited.

Here, we report an expression system for recombinant production of N-terminally acetylated SOD1 in *E. coli.* To understand the effect of N-terminal acertylation of SOD1 on misfolding and self-assembly the modified protein is characterized in terms of apo-state structure and stability and the propensity for amyloid formation and compared to SOD1 with a free N-terminus. The experiments show that amyloid formation is accelerated by N-terminal acetylation both by shortening of the lag-phase and increase of the elongation rate. At the same time, the SOD1 apo structure and stability are largely unaltered by the modification, and no changes are observable in the morphology of fibrils.

## Results

### Production of recombinant N-terminal acetylated superoxide dismutase 1

In general, bacterial cells lack the apparatus for N-terminal acetylation of proteins. A consequence of this is that heterologous expressed proteins in *E. coli* have a free N-terminus. In contrast, many eukaryotic proteins are acetylated at the N-terminus (figure 1). The two first residues in human SOD1 are methionine and alanine. The methionine is cleaved off during translation, as expected when the second residue is alanine, also in *E. coli*. To produce SOD1 acetylated at the N-terminus, we developed an expression system where SOD1 is co-expressed with the NAT A catalytic subunit NAA10 and ribosome anchoring subunit NAA15 from *Schizosaccharomyces pombe*. The two enzyme subunits have previously been reported to express well and be active in *E. coli*(14). The NAT A genes were expressed in the pCDFDuet vector from Novagen encoding for spectinomycin resistance and human SOD1 with the six mutations C6A, F50E, G51E, C57S, C111A, C146S referred to in the following as pwtSOD1^Δcys^ were express in the pET17b vector carrying a gene for ampicillin resistance as previously described (15). This SOD1 variant was chosen to achieve SOD1 in the monomeric state, limiting the formation of the native dimer (F50E and G51E) and avoiding the formation of disulfide bonds by mutation of the four cysteine residues (C6A, C57S, C111A, and C146S) (16, 17). The *E. coli* BL21DE(3) pLysS (chloramphenicol resistance) strain was co-transformed with the two plasmids. Protein expression was performed either in rich LB medium or in high salt M9 minimal medium supplemented with glucose and ^15^N-NH_4_Cl under selection by ampicillin, chloramphenicol, and spectinomycin to produce uniformly ^15^N-labelled protein for NMR. Protein expression was induced by the addition of isopropyl β-d-1-thiogalactopyranoside (IPTG). N-terminal acetylated pwtSOD1^Δcys^ (N_term_-acetyl-pwtSOD1^Δcys^) was purified essentially as described (15). The presence of the N-terminal acetylation modification was confirmed by mass spectrometry MALDI-TOF and TIMS-TOF MS/MS after proteolytic cleavage by trypsin and AspN. Anionic exchange chromatography and mass-spectrometry show that N-terminal acetylation is nearly complete after co-expression with Naa10 and Naa15 as pwtSOD1^Δcys^ with free N-terminus is not detected.

### N-terminal acetylation does not affect SOD1 secondary structure and thermal stability of the apo-state

SOD1 has an immunoglobulin fold with eight anti-parallel beta-strands arranged in a beta-sandwich with anti-parallel strands connected by loops (figure 1c). The secondary structure of N_term_-acetyl-pwtSOD1^Δcys^ was probed by circular dichroism (CD) and nuclear magnetic resonance (figure 2). There are no significant changes of the far-UV CD spectrum of N_term_-acetyl-pwtSOD1^Δcys^ compared to the apo protein with a free N-terminal (pwtSOD1^Δcys^). Upon heating changes to the far-UV CD spectrum indicate unfolding of secondary structure elements within the protein (figure 2a).

**Figure 2.**
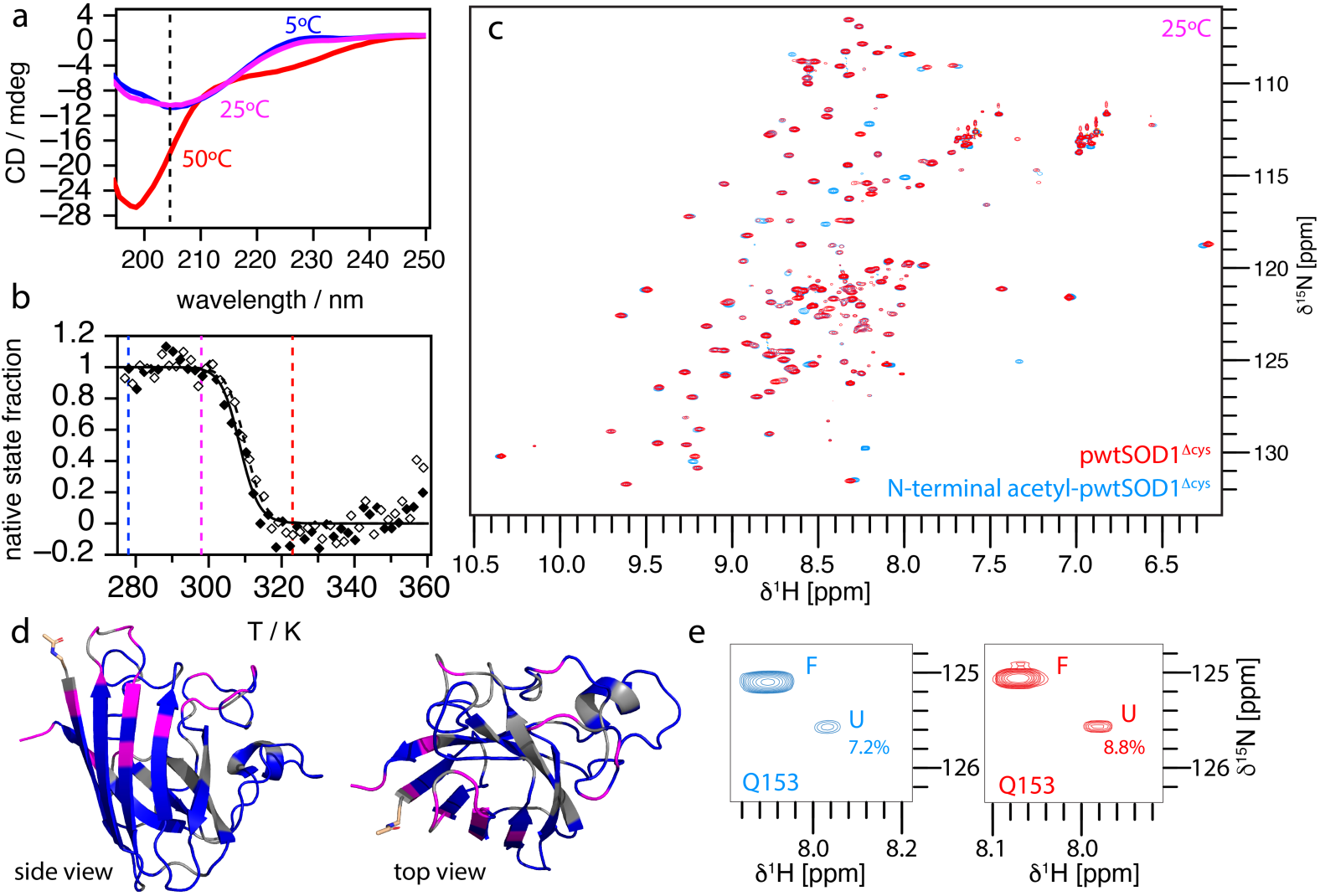
Effect of N-terminal acetylation on thermal stability of pwtSOD1^Δcys^. a) Circular dichroism (CD) spectra of 10 µM N_term_-acetyl-pwtSOD1^Δcys^ at different temperatures in 10 mM sodium phosphate pH 7.0. No significant changes on the far-UV CD-spectra are observed upon acetylation when compared to pwtSOD1^Δcys^. b) The thermal stability was determined by recording of the CD signal at 205 nm heating the sample from 5°C to 85°C in 1°C steps with 60 second equilibration time at each temperature before measurement. c) Overlay of ^1^H-^15^N-HSQC spectra of 0.65 mM ^15^N-labelled N_term_-acetyl-pwtSOD1^Δcys^ (blue) and of 0.5 mM ^15^N-labelled pwtSOD1^Δcys^ (red) recorded at 25°C in 10 mM sodium phosphate pH 7.0. d) Side view and top view of the SOD1 monomer (2xjl) in cartoon representation showing residues that display chemical shift perturbations upon N-terminal acetylation in magenta. Residues missing in the pwtSOD1^Δcys^ spectra based on assignments deposited under BMRB entry 15711 (21) are shown in gray. e) Close view on the ^1^H-^15^N-HSQC spectra from c) showing the cross peaks originating from the C-terminus residue Gln153 in the folded (F) and unfolded (U) states.

The effect of N-terminal acetylation on thermal stability was investigated by following CD signal changes at 205 nm as a function of temperature. This choice was made based on the large change in CD signal with temperature for pwtSOD1^Δcys^ at that wavelength (figure 2a). The thermal unfolding of N_term_-acetyl-pwtSOD1^Δcys^ and pwtSOD1^Δcys^ were found to be cooperative and reversible (figure 2b) indicating a two-state system in accordance with previous reports of the SOD1 unfolding (16, 18). The thermal transition data was fitted to equation 1 neglecting any differences in the heat capacity between the folded state and unfolded state (Δ*C_p_* = 0). This assumption is supported by reports of Δ*C_p_* for unfolding of the SOD1 apo-state to be low due to the dynamic nature of the folded apo-state (19). The mid-point of the transition for pwtSOD1^Δcys^ was 309.6 ± 0.7 K and 308.0 ± 0.9 K for N_term_-acetyl-pwtSOD1^Δcys^. These results demonstrate that the stability of the apo state of the two variants is, within error, the same (figure 2b).

The folding status of the SOD1 variants was further confirmed by recording backbone amide ^1^H-^15^N HSQC spectra of both variants (N_term_-acetyl-pwtSOD1^Δcys^ and pwtSOD1^Δcys^) at 25°C (figure 2c). The ^1^H-dimension of both spectra was well dispersed and in accordance with previous spectra of folded SOD1 (20, 21). To identify residues that are affected by the acetylation chemical shift perturbations were assessed. Most peaks have unaltered chemical shifts upon N-terminal acetylation when compared to the protein with free N-terminus. Due to the lack of chemical shift assignments of N_term_-acetyl-pwtSOD1^Δcys^ variant, a detailed analysis of chemical shift perturbations is difficult. Instead peaks that either move or are only found in the spectrum of pwtSOD1^Δcys^ were identified and plotted on the structure SOD1 (figure 2d). This analysis identifies residues that are affected by the modification. It shows that residues at the N-terminal and proximal to the N-terminus in loop II display significant chemical shift perturbations. In addition, residues forming a belt across the β-sheet formed by strands β1, β2, β3, and β6 also show perturbations. Based on previous analysis of SOD1 variants with a free N-terminus (15, 20, 22) it is evident that a subpopulation of both variants is unfolded at 25°C, the temperature where the spectra were recorded (figure 2c and 2e). The unfolded fraction may be estimated from the ratio of the volumes of the C-terminus cross peaks (residue Q153) originating from the folded (F) and unfolded (U) states (22). This analysis gives similar U/F ratios for both variants and in accordance with the ratio at 25 °C obtained from CD spectroscopy. Collectively the CD spectroscopy and NMR analysis shows that N_term_-acetyl-pwtSOD1^Δcys^ is folded into a structure that resembles the fold and stability of pwtSOD1^Δcys^.

### Amyloid formation is accelerated by N-terminal acetylation

To assess the impact of N-terminal acetylation on amyloid formation, the self-assembly kinetics were monitored in a coupled assay by fluorescence of the amyloid-specific dye thioflavin T (ThT). ThT fluorescence was chosen as a reporter to monitor reaction progress due to its high sensitivity and signal-to-noise ratio (23). This is particularly important when covering larger protein concentration spans. Aggregation kinetics were measured as a function of the monomer concentration of N_term_-acetyl-pwtSOD1^Δcys^ ranging from 2.2 µM to 140 µM and compared to the variant with a free N-terminal amino group at 10 mM sodium phosphate pH 7.0, 0.5 M NaCl, 75 µM ThT at 37 °C with orbital shaking (Fig. 3). Fibril formation as monitored by changes in ThT-fluorescence of the N_term_-acetyl-pwtSOD1^Δcys^ variant shows a clear dependence on protein concentration with sigmoidal curve shapes. The reactions are accelerated compared to pwtSOD1^Δcys^ (figure 3). Curves from reactions with concentrations of 17.5 µM N_term_-acetyl-pwtSOD1^Δcys^ or higher all show steep transitions within the time frame of the experiment. This is like curves reported for other fibril-forming proteins and indicates the involvement of secondary processes during fibril formation (24). Increasing the concentration of N_term_-acetyl-pwtSOD1^Δcys^ leads to a shortening of the observed lag phase (figure 3a, 3b and 3d). The lag-phase length dependence on initial monomer concentration follows a power-law for both SOD1 variants, with a stronger dependence on monomer concentration for the N-terminal acetylated variant (figure 3d). Fitting of a power law to the data yields an exponent of −0.4 ± 0.1 for the N-terminal acetylated variant and −0.1 ± 0.1 for the variant with a free N-terminus.

**Figure 3.**
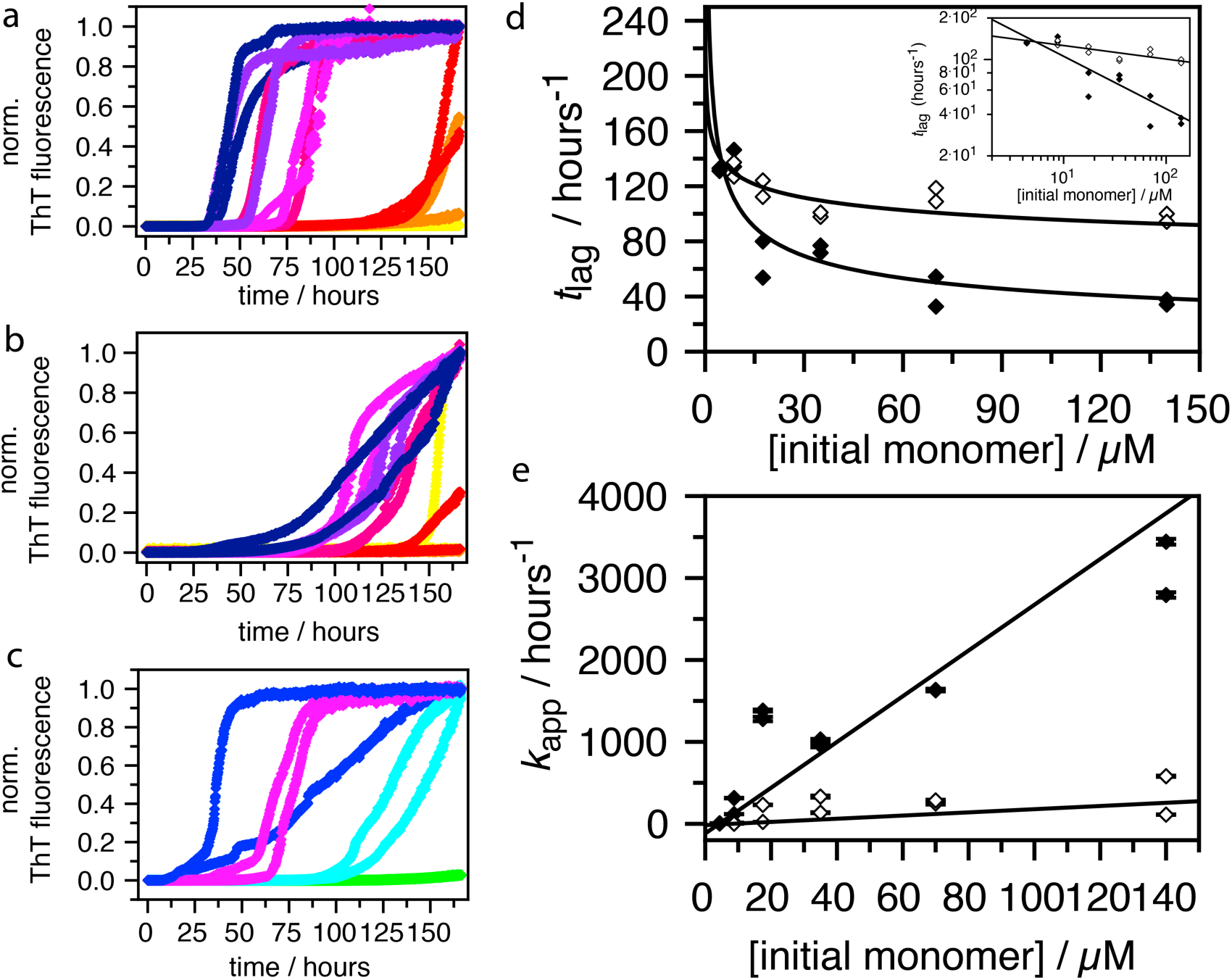
Effect of N-terminal acetylation on SOD1 amyloid formation kinetics. Kinetic traces based on measurements of ThT fluorescence in assays with increasing concentrations of **a)** N_term_-acetyl-pwtSOD1^Δcys^ 2.2 µM (yellow), 4.4 µM (orange), 8.8 µM (red), 17.5 µM (pink), 35.0 µM (magenta), 70.0 µM (purple) and 140.0 µM (blue). All reactions were done at 37°C, 10 mM sodium phosphate pH 7.0, 0.5 M NaCl, 75 µM ThT with orbital shaking and **b)** pwtSOD1^Δcys^ 2.2 µM (yellow), 4.4 µM (orange), 8.8 µM (red), 17.5 µM (pink), 35.0 µM (magenta), 70.0 µM (purple) and 140.0 µM (blue). All reactions were done at 37°C, 10 mM sodium phosphate pH 7.0, 0.5 M NaCl, 75 µM ThT with orbital shaking. **c)** 35 µM N_term_-acetyl-pwtSOD1^Δcys^ in 10 mM sodium phosphate pH 7.0 and 75 µM ThT with different concentrations of NaCl: 1 M (blue),0.5 M (magenta), 0.25 M (cyan) and 0.125 M (green) at 37°C with orbital shaking. **d)** Plot of the time t_lag_ as function of initial monomer concentration, N_term_-acetyl-pwtSOD1^Δcys^ (filled diamonds) and pwtSOD1^Δcys^ (open diamonds). Functions are fit of equation 3 to the data points. Insert: same data and functions shown in a double logarithmic plot. e) Plot of the apparent rate constant obtained from the maximum slope of the kinetic trances as function of initial monomer concentration, N_term_-acetyl-pwtSOD1^Δcys^ (filled diamonds) and pwtSOD1^Δcys^ (open diamonds). Functions are linear fits to the data points resulting in second order rate constants of elongation of (27.9 ± 0.1)10^6^ M^−1^hours^−1^ for N_term_-acetyl-pwtSOD1^Δcys^ and (1.9 ± 0.1)10^6^ M^−1^hours^−1^ for pwtSOD1^Δcys^.

Estimates of the effect on fibril elongation can be obtained either from sigmoidal fitting to the data or from linear fitting of the elongation phase to determine the maximum reaction rate (25). From this analysis, an apparent rate constant of elongation can be found from the maximum slope. This was done for N_term_-acetyl-pwtSOD1^Δcys^ and pwtSOD1^Δcys^ to compare the effect of N-terminal acetylation on fibril elongation. Through plotting of the apparent rate constant as a function of the initial monomer concentration, estimated second-order rate constants for fibril elongation were obtained yielding *k_el_* = (27.9 ± 0.1)10^6^ M^−1^hours^−1^ for N_term_-acetyl-pwtSOD1^Δcys^ and *k_el_* = (1.9 ± 0.1)10^6^ M^−1^hours^−1^ for pwtSOD1^Δcys^ (figure 3e). This analysis shows that the second order elongation rate constant is increased in the order of 10-fold by N-terminal acetylation.

The self-assembly reactions were done under conditions of high ionic strength obtained by having 0.5 M NaCl in the reactions. Under these conditions Coulomb interactions are screened which may reduce repulsion of similar charged molecules. Increasing the ionic strength shortens the lag phase and accelerate elongation rates of N_term_-acetyl-pwtSOD1^Δcys^ (figure 3c). Similar effects are observed for pwtSOD1^Δcys^. Amyloid formation of both variants is accelerated by salt screening, but screening of Coulomb interactions in the pwtSOD1^Δcys^ self-assembly reaction is not enough to obtain the same fibrillation rates as for the N_term_-acetyl-pwtSOD1^Δcys^ variant indicating that the effect of acetylation cannot be explained alone by changes in electrostatic interactions originating from local removal of a positive charge upon acetylation of the N-terminus.

Collectively, the kinetic analysis shows that N-terminal acetylation accelerates self-assembly by modulation of both primary nucleation and elongation. However, more investigations are needed to identify the underlying molecular mechanism behind the increased aggregation propensity of N_term_-acetyl-pwtSOD1^Δcys^.

### SOD1 fibril morphology is insensitive to N-terminal acetylation

The effect of N-terminal acetylation on fibril morphology was investigated by cryogenic electron microscopy imaging of the formed fibrils after two weeks of incubation at 37°C under ThT-assay conditions with 17.5 µM (35 µM) N_term_-acetyl-pwtSOD1^Δcys^ (pwtSOD1^Δcys^) in 10 mM sodium phosphate pH 7.0, 0.5 M NaCl and 75 µM ThT (figure 4). Under these conditions, Coulomb interactions are screened by the high concentration of salt ions. At the time-point for fibril extraction, there were many more fibrils in the N-acetylated sample than in the sample with free N-terminus reflecting the difference in reaction kinetics. Despite this difference in fibril concentration between variants, specimens of thin-layer vitrified samples were prepared for cryogenic electron microscopy as described in the method section. Images collected on different grids, and on different parts of the same grid were recorded. Qualitatively, the SOD1 fibrils can be described as long (several µm in length) and unbranched with a low degree of twisting. Fibrils are found twisted around each other forming long fibrillar bunches of seemingly two to three fibrils. The resolution of these images does not allow for more detailed quantification of fibril fine structures, twisting, and topologies. Fibril morphology was further evaluated using the program ImageJ (26). The fibril cross-section width was measured multiple times in different images and the average length was found by fitting a Gaussian function to the measured widths (figure 4b and d). The observed average fibril widths were similar for the two variants with N_term_-acetyl-pwtSOD1^Δcys^ fibrils having an average width of (7.4 ± 0.1) nm with a standard deviation of (1.2 ± 0.1) nm compared to (7.0 ± 0.1) nm with a standard deviation of (1.0 ± 0.1) nm on average for fibrils of the variant with free N-terminus. By comparison to the published cryogenic electron microscopy structure of wild type SOD1 (27) (figure 4e) the observed fibril morphology is consistent with having one monomer in each fibril layer. Collectively, the analysis shows that N-terminal acetylation of pwtSOD^Δcys^ has little effect on the observed fibril morphology when evaluated by cryogenic electron microscopy imaging.

**Figure 4.**
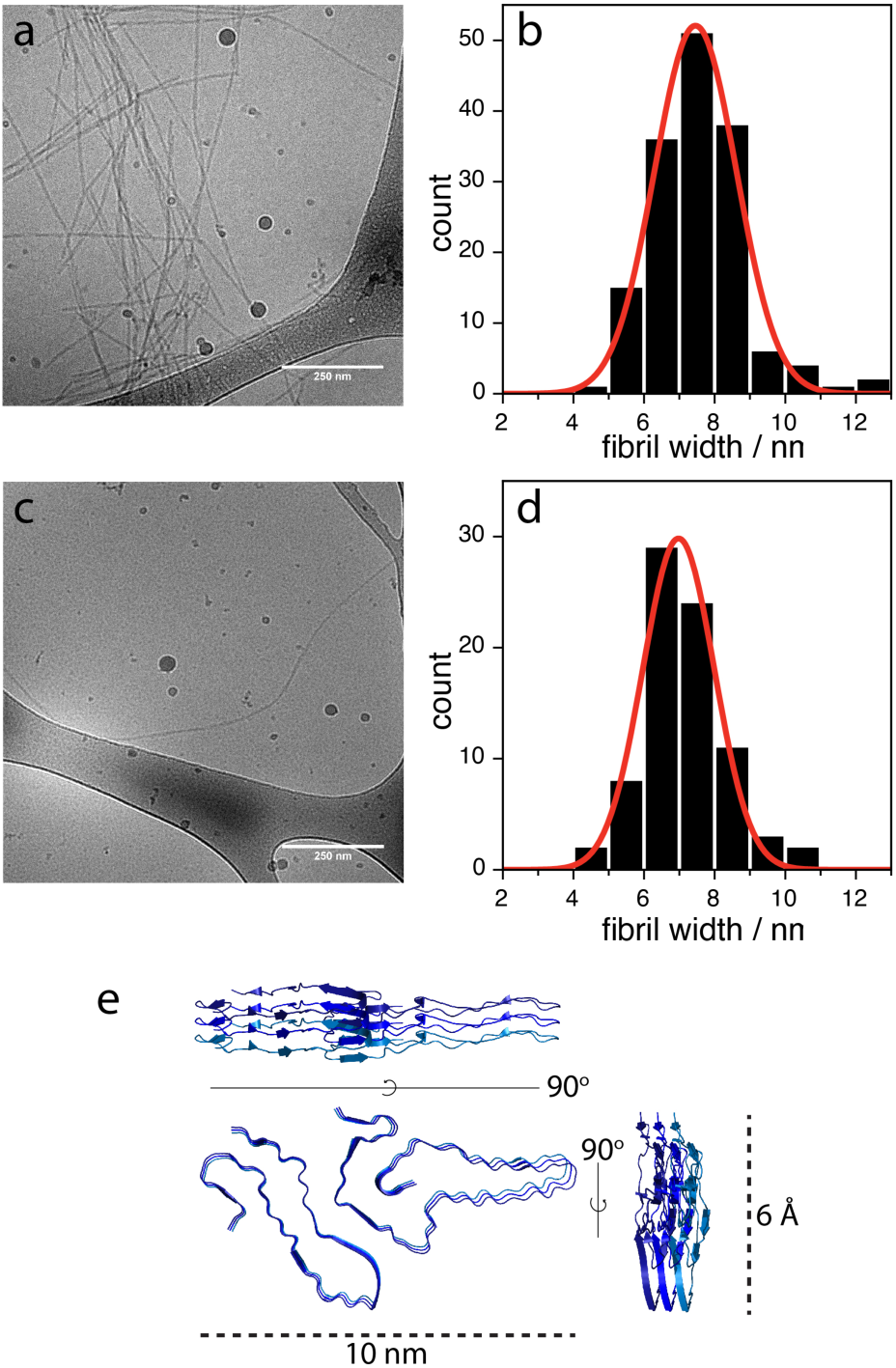
Effect of N-terminal acetylation on fibril morphology. cryogenic electron microscopy micrograph of a) 35 µM N_term_-acetyl-pwtSOD1^Δcys^ and c) 35 µM pwtSOD1^Δcys^ after two weeks of incubation at 37°C with orbital shaking in 10 mM sodium phosphate pH 7.0, 75 µM ThT and 0.5 M NaCl. Histogram of measured fibril widths from images b) N_term_-acetyl-pwtSOD1^Δcys^ and d) pwtSOD1^Δcys^. Red curves are fits of a gaussian function (equation 4) to the width distributions. e) cartoon representation of three chains each constituting a layer in the amyloid structure of human wild type SOD1 (PDB entry 7vzf)(27). Dimensions of the fibril structure are shown for comparison to the measured fibril widths.

## Discussion

SOD1 is a cytosolic enzyme with extraordinary stability in its dimeric metal ion-bound holo-state (16). In contrast, the stability of the monomeric apo-state is significantly reduced. Several studies have identified metal depletion and/or reduction of the intramolecular disulfide bond in SOD1 to lead to decreased stability and accelerated amyloid formation (16, 17, 28). SOD1 apo-state folding can be described by a two-state mechanism followed by metal ion binding and dimerization into the holo-state (16). The mechanisms of protein amyloid formation are, in general complex with the possibilities of multiple parallel pathways leading from monomer to the polymeric amyloid fibril (1, 24). Chemical denaturant studies of the aggregation propensity of SOD1 have suggested that SOD1 may aggregate from the partly folded apo-state or from the fully unfolded state (17, 18, 21). Making more than one route into the amyloid pathway likely depending on reaction conditions. Studies of the slow µs-ms dynamics of the SOD1 apo state show that SOD1 builds transient non-native oligomers (21, 29) likely to be obligatory for amyloid formation. Studies of the amyloid-forming reaction with translational diffusion NMR and cross-linking mass-spectrometry (15, 30) and atomic force microscopy (31, 32) have demonstrated SOD1 to form soluble oligomers in the size range of trimers to tetramers during aggregation and amyloid formation. The formed oligomers are reported to be *off*-pathway for amyloid formation and found to be toxic to cells (31, 33).

N-terminal acetylation leads to a change in charge from positive to neutral, leading to an overall more negative net charge of SOD1, which is negatively charged at physiologically pH. This means that a simple electrostatic mechanism cannot account for the observed acceleration of amyloid formation kinetics upon acetylation as acetylation then should be expected to retard self-assembly by increased electrostatic repulsion as previously reported for unspecific acetylation of lysine side chains in SOD1 (34). The present findings show that N-terminal acetylation accelerates amyloid formation in the presence of high concentrations of salt ions i.e. under conditions where coulomb interactions are screened and intermolecular repulsion dampened. This suggests that the effect of N-terminal acetylation is decoupled from pure electrostatic effects originating from having a charged N-terminus in non acetylated SOD1.

The finding that the apo-state stability is not changed by acetylation of the N-terminus while the amyloid formation kinetics is accelerated is consistent with the behavior of a subset of the more than 200 single-point mutations in SOD1 associated with ALS that result in gain-of-toxicity and disease, but do not destabilize the apo-state (35, 36). Most of these non-destabilizing ALS mutations are found on the surface of SOD1. Given the complex multi-step reaction pathway leading to amyloid formation, it is expected that modifications may affect kinetics without interference with the structure and stability of the starting state. A comparison of the width of the fibrils formed by both SOD1 variants with the single particle cryoEM structure of wtSOD1 (27) indicates that only one SOD1 chain is found in each fibril layer. This may be interpreted as fibril elongation proceeds as a bimolecular reaction between fibril-ends and monomeric SOD1 however, this mechanism is not proven.

A possible mechanistic explanation is that N-terminal acetylation affects either minor invisible states or the formation of *off*-pathway soluble oligomers. The direct consequence of destabilization of *off*-pathway oligomers would be to increase the effective concentration of reactive monomeric species. This would increase the rate of primary nucleation with a shortened lag phase while increasing the rate of elongation by increasing the concentration of one of the species involved in the elongation step. Alternatively, acetylation may expose and populate transient structures that lead to increased primary nucleation and fibril elongation. However, further studies are needed to elucidate the underlying molecular mechanism that leads to the acceleration of SOD1 amyloid formation by N-terminal acetylation. It also remains to be investigated whether N-terminal acetylated SOD1 is more or less toxic compared to SOD1 with a free N-terminus.

The present study clearly points to the importance of investigating the effect of co- and posttranslational modifications on protein self-assembly and misfolding as even a minor modification is shown to have large impact on amyloid formation kinetics. Some general effects on e.g. protein stability and cellular half-life may exist for individual modifications, e.g. N-terminal acetylation may act as an N-degron to protect against degradation via the ubiquitin-proteasome system (12). However, specific PTMs are expected to act with different effect depending on the modified protein, as is the case for N-terminal acetylation that was previously found to slow down amyloid formation of α-synuclein (37). Understanding the effect of PTMs on amyloid formation is key for further understanding of the molecular events in protein misfolding diseases and their complex phenotypes.

## Material and Methods

### Protein expression and purification

N-acetyl- pwtSOD1^Δcys^ was produced by co-expression of human SOD1^Δcys^ C6A, F50E, G51E, C57S, C111A, C146S referred to here as pwtSOD1^Δcys^ in a pET17b vector carrying a gene for ampicillin resistance as described in (15) with pCDFDuet vector (Novagen) encoding for spectinomycin resistance harboring ribosome anchoring subunit NAA15 from *Schizosaccharomyces pompe* (residue 1 – 729) in multiple cloning site I and catalytic subunit NAA10 from *Schizosaccharomyces pompe* (residue 1 – 177) in multiple cloning site II (synthesized by GenScript). The *E. coli* BL21DE(3) pLysS (chloramphenicol resistance) strain was co-transformed with the two plasmids. Protein expression was performed either in rich LB medium or in high salt M9 minimal medium supplemented with glucose and ^15^N-NH_4_Cl under selection by ampicillin, chloramphenicol and spectinomycin to produce uniformly ^15^N-labelled protein for NMR. Protein expression was induced by addition of isopropyl β-d-1-thiogalactopyranoside (IPTG) to 0.8 mM and at the same time CuSO_4_ and ZnCl_2_ were added to a final concentration on 60 µM, and the cultures were left for O.N. expression at RT under shaking conditions as described (15, 21). N-terminal acetylated pwtSOD1^Δcys^ (N_term_-acetyl-pwtSOD1^Δcys^) was purified essentially as described (15, 16). Protein concentrations were determined by absorbance at 280 nm using a NanoDrop 2000/2000c spectrophotometer (Thermo Scientific) and calculated using an extinction coefficient of ε_280nm_ = 5500 M^−1^cm^−1^.

### Mass spectrometry

The purified proteins were analysed by intact mass spectrometry by recording of MALDI-TOF/TOF MS and MSMS spectra using a Autoflex Speed MALDI TOF/TOF mass spectrometer (Bruker Daltonics, Bremen, Germany) in positive reflector mode, see SI for details. All spectra were externally calibrated using Peptide calibration standard II (Bruker Daltonics).

N-terminal acetylation was confirmed by in-solution proteolytic digest of the purified protein variants by trypsin (Promega, Madison, WI, USA) and AspN (Promega, Madison, WI, USA) followed by purification of the peptide fragments by reverse-phase HPLC. First the peptides were cleaned up on a C18 micro column before injected onto to an ultra-high pressure nanoflow chromatography system (nanoElute, Bruker Daltonics). The peptides were loaded onto an Acclaim PepMap C18 (5 mm, 300 μm id, 5 μm particle diameter, 100 Å pore size) trap column (Thermo Fisher Scientific) and separated on a Bruker Pepsep Ten C18 (75 µm × 10 cm, 1.9 µm particle size) analytical column (Bruker Daltonics). Peptides were analysed on a quadrupole time-of-flight mass spectrometer (timsTOF Pro, Bruker Daltonics), via a nano electrospray ion source (Captive Spray Source, Bruker Daltonics) in positive mode, controlled by the OtofControl 5.1 software (Bruker Daltonics). For details see SI.

#### Data analysis

Raw data from the LC-MSMS analysis were processed using Mascot Distiller (version 2.8) and searched against an in-house database containing the pwtSOD1^Dcys^ protein sequence using the settings; precursor ion tolerance 8 ppm, MS/MS fragment mass tolerance 0.015 Da, trypsin and Asp-N as protease, 1 missed cleavage site, oxidation (M) and acetylation (N-term) as variable modification.

### Circular dichroism

The initial degree of secondary structure under different conditions were propped by circular dichroism using a J-815 spectropolarimeter (Jasco, Mary’s Court Easton, MD, USA) equipped with a peltier element for precise temperature control. Quarts cuvettes with 1 mm light path (Hellma Analytics, Müllheim, Germany) were used for all measurements. Measurements of monomeric N-acetyl-pwtSOD1^Δcys^ and pwtSOD1^Δcys^ were performed in 10 mM sodium phosphate pH 7.0 with varying concentration of sodium chloride (0 M, 0.15 M and 0.5 M) and at different temperatures (5°C, 25°C, 50°C) with protein concentrations of either 10 µM or 25 µM.

Thermal stability measurements were performed with a heat increase of 0.5°C/min and a 60 s equilibration time at each temperature (steps of 1°C). The CD signal at 205 nm was recorded with a 2 nm bandwidth and an 8 s digital integration time between 5°C and 85°C in steps of 1°C. Both forward (heating) and reverse (cooling) was performed to check for folding reversibility. Thermal transition data were analysed by fitting of the Gibbs-Helmholtz equation (Eq. 1) to the data assuming two state reversible folding/unfolding of N-acetyl-pwtSOD1^Δcys^:

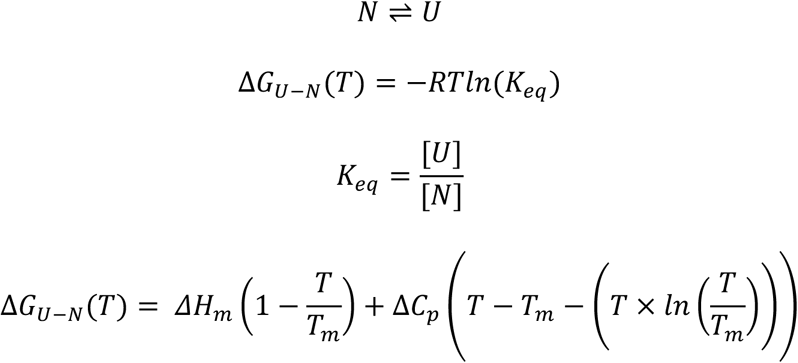

Where *T_m_* is the temperature where *ΔG_D-N_(T)* = 0 and Δ*H_m_* is the enthalpy at *T_m_*. *ΔC_p_* is the difference in heat capacity between the folded (N) and unfolded (U) state and assumed to be invariant with temperature in the temperature range studied.

Assuming similar heat capacities of the folded and unfolded states i.e. Δ*C_p_* equal to zero equation 1 reduces to:

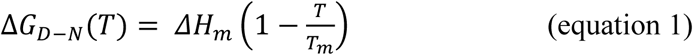

### NMR spectroscopy

Two dimensional ^1^H,^15^N-HSQC spectra of uniformly ^15^N-labelled 0.65 mM N_term_-acetyl-pwtSOD1^Δcys^ and 0.5 mM pwtSOD1^Δcys^ in 10 mM Na-PO_4_, pH 7.0, 10% D_2_O were recorded at 25°C in 5 mm precision tubes (Wilmad 535-PP-7) on a Bruker Avance Neo 14.1 Tesla 4-channel spectrometer equipped with a QCI(P) 5mm Z-gradient CryoProbe (1H-19F)/13C/15N/31P/2H. Data were processed in NMRpipe (38) and analysed in ccpnmr V3 analysis (39). Chemical shift assignments of pwtSOD1 recorded at 25°C from the BMRB databank entry ID 15711 was used in combination with previous assignment transfer to pwtSOD1^Δcys^ at 37°C (15, 20, 21). Peak integration were performed in PINT (40).

### Thioflavin T coupled amyloid formation assay

Amyloid formation was probed in a ThT-coupled assay. The reaction mixtures contained 75 µM ThT, 10 mM sodium phosphate pH 7.0 and either 0.5 M NaCl and varying concentration of N-acetyl pwtSOD1^Δcys^ (2.2, 4.4, 8.8, 17.5, 35.0, 70.0, 140.0 µM) or 35 µM N_term_-acetyl-pwtSOD1^Δcys^ and varying concentration of sodium chloride (0, 0.0625, 0.125, 0.25, 0.5, 1 M). The assay was performed with a sample volume of 100 µl in 96-well Corning 3881 half area plates sealed with SealPlate® plastic film (Excel Scientific) at 37°C in a FLUOstar Omega plate reader from Thermo Fisher using bottom reading with a 448 nm excitation filter and a 480 nm emission filter with 20 second orbital shaking 300 rpm before each detection round (600 second cycle periods). The time dependent fluorescence intensity data was fitted to equation 2 (23):

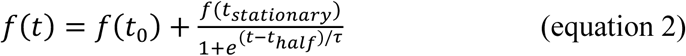

Where *f(t_0_)* is the initial fluorescence signal and *f(t_stationary_)* is the fluorescence signal at the final plateau, *t_half_* is the time point halfway through the elongation phase and τ is the time constant for elongation equal to the reciprocal of *k_app_* the apparent rate constant for elongation. Reliable fitting of a sigmoidal equation (equation 2) to the experimental data was difficult for the reactions displaying slow aggregation kinetics, mainly due to that the reactions did not reach the stationary phase. Therefore, *t_half_* was also found for all kinetic traces by fitting of a spline function to the data and determination of the maximum of its first derivative and compared to the values obtained from fitting to equation 2. Apparent rate constants of elongation were also estimated from the slope of linear fits to the elongation phase data points. Times of *t_lag_* were determined as the time point where the linear fit function to the elongation phase intercept with a linear fit function to the lag-phase data. Linear correlation was found for *t_lag_* and *t_half_* (correlation coefficient = 0.96).

Lag-phase length dependence of initial monomer concentration was fitted to a power law:

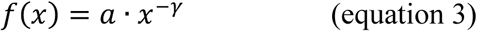

### Cryogenic electron microscopy

Samples were taken from the ThT-reactions with 35 µM N_term_-acetyl-pwtSOD1^Δcys^ (pwtSOD1^Δcys^) after 167 hours of incubation where the N_term_-acetyl-pwtSOD1^Δcys^ had reached a plateau in ThT fluorescence and prepared for cryogenic electron microscopy imaging. Specimens were prepared as thin liquid films on glow discharged lacey carbon-coated copper grids (Ted Pella), blotted with filter paper and plunged into liquid ethane (−180°C) in a controlled environment using a Vitrobot. Vitrified specimens were stored under liquid nitrogen until measured. Specimens were transferred to the JEOL 200kV electron microscope using a Fischione model 2550 cryo-transfer tomography holder. Zero-loss images were recorded digitally with a TVIPS F416 camera using SerialEM under low-dose conditions with a 30 eV energy selecting slit.

Image analysis was performed in Image J (26). The width of fibrils was determined by averaging multiple measurements carried out on different fibrils and on different images as well as multiple measurements of the same fibril along the fibril axis. A total of 154 (81) measurements of fibril width of N_term_-acetyl-pwtSOD1^Δcys^ (pwtSOD1^Δcys^) were included in the analysis. The average fibril widths were determined by grouping the measurements into bins of 1 nm steps and fitting to a gaussian function:

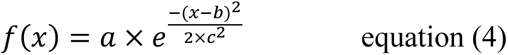

where a is the maximum hight (count number), b is the position of the maximum (nm) and c is the standard deviation (nm).

## Acknowledgement

This work was funded by a grant from Swedish research council (202004888), The Crafoord Foundation (20230966), Per-Eric and Ulla Schyberg’s Foundation, Åke Wiberg’s Foundation (M23-0143), The Danish council for independent research (4002–00258) and Royal Physiographic Society of Lund.

Sara Linse is thanked for the kind access to the FLUOROstar Omega plate reader. The excellent technical support from Crispin Hetherington with cryoEM imaging, Katja Bernfur with mass spectrometry and Zoltan Takacs with NMR spectroscopy is acknowledged. NMR spectra were recorded at the NMR Center at the Department of Chemistry, Lund University.

## Abbreviations

SOD1: superoxide dismutase 1
ALS: amyotrophic lateral sclerosis
N_term_-acetyl-pwtSOD1^Δcys^: N-terminal acetylated SOD1 C6A/F50E/G51E/C57S/C111A/C146S
pwtSOD1^Δcys^: SOD1 C6A/F50E/G51E/C57S/C111A/C146S
NAT A: N-terminal acetyl transferase A
NMR: nuclear magnetic resonance
CD: circular dichroism
MS: mass-spectrometry

